# Immune profiling of cord blood after prolonged rupture of membranes

**DOI:** 10.1101/444216

**Authors:** Evdokia Tsaliki, Carolin T Turner, Cristina Venturini, Christy Kam, Angela Strang, Sarah Bailey, Alison Rodgers, Adam P Levine, Benjamin M Chain, Nandi Simpson, Eleanor M. Riley, Nigel Field, Peter Brocklehurst, Mahdad Noursadeghi

## Abstract

We hypothesised that foetal immune responses to an infectious challenge may be detected by genome-wide transcriptional profiling of cord blood. In order to test this hypothesis, we sought to identify transcriptomic changes in post-natal cord blood samples following prolonged pre-labour rupture of membranes (PROM) as a surrogate for increased risk of infection. By comparison to controls we found increased levels of blood transcripts in a subset of prolonged PROM cases, significantly enriched for innate immune system signalling pathways. These changes were idiosyncratic, suggesting qualitative and quantitative variation in foetal immune responses which may reflect differences in exposure and/or in host genetics. Our data support the view that PROM represents an infection risk to the foetus. In addition, we propose that cord blood transcriptional profiling offers exciting opportunities to identify immune correlates of clinical outcome following potential in utero exposures to infection. These may be used to elucidate the mechanisms of immunological protection and pathology in the foetus and identify biomarkers to stratify the risk of adverse outcomes.

## Introduction

Immune responses to antenatal infections are poorly characterised. Targeted proteomics and mass cytometry have recently revealed inflammatory responses in the cord blood of preterm infants, pointing to immune responses associated with the underlying pathology (1). However, this approach is not easily scalable. In recent years there has been substantial interest in studying the immunology of infectious disease in children and adults, using genome-wide transcriptional profiling of whole blood (2). This approach is based on the premise that peripheral blood leukocytes transit through immunologically active sites and thereby reflect active immunological pathways within their transcriptome. It takes advantage of readily accessible and scalable technologies that minimise technical confounding by stabilising RNA as blood is drawn into proprietary tubes and depletion of globin mRNA in order to increase the sensitivity for changes in the transcriptome of blood leukocytes. In the present study, we sought to apply this approach to investigating antenatal immune responses by transcriptional profiling of unfractionated cord blood.

Prolonged pre-labour rupture of membranes (PROM) is thought to incur higher risk of ascending intrauterine infection, leading to expedited delivery and/or the empirical use of perinatal antimicrobial drugs (3, 4). Therefore, we hypothesised that transcriptional profiling of cord blood obtained at the time of delivery, will reflect foetal immune responses to ascending infection amongst prolonged PROM cases compared to controls. This would provide proof of concept for the application of cord blood transcriptomics to investigate foetal immune responses in order to explore immune correlates of protection or pathology following *in utero* exposure to an infectious insult, and potentially to identify biomarkers of differential clinical outcomes following such an exposure.

## Results and discussion

We collected cord blood samples from 20 consecutive neonates with prolonged PROM of greater than 24 hours before delivery and from 81 controls, represented by consecutive neonates with rupture of membranes less than 24 hours before delivery (Table 1). The mean interval time between rupture of membranes and delivery in PROM cases was 40 hours and in controls was 4.5 hours. Gestational age, mode of delivery, birth weight, APGAR score and in-hospital length of stay were not significantly different between cases and controls.

**Table 1:** Clinical metadata for PROM cases and controls.

|  |  | PROM | Control |
| --- | --- | --- | --- |
| **Number** |  | 20 | 80 |
| **Mean gestation (range)** |  | 40 (37-42) | 39 (36-42) |
| **Mean ROM duration (range)** |  | 41.1(24-61) | 4.5 (0-22) |
| **% Mode of delivery** | **Spontaneous vaginal birth** | 45 | 42 |
|  | **Instrumental vaginal birth** | 15 | 9 |
|  | **Caesarean section** | 40 | 49 |
| **Mean birth weight in kg (range)** |  | 3.4 (2.6-4.4) | 3.3 (2.4-4.5) |
| **Mean APGAR<sub>5</sub> score (range)** |  | 9.4 (8-10) | 10 (8-10) |
| **Mean hospital LOS (range)** |  | 1.7 (0-5) | 2 (0-8) |

Abbreviations: ROM (rupture of membranes); APGAR<sub>5</sub> (Appearance, Pulse, Grimace, Activity, Respiration score at 5 minutes; LOS (length of stay).

In principal component analysis of genome-wide cord blood transcriptomes, samples from PROM cases and controls clustered together (Figure 1A). Grouping all PROM cases together, we found statistically significant enrichment of 27 protein-coding transcripts. In this analysis we did not use multiple testing corrections in order to minimise the likelihood of type 2 errors in the identification of differentially expressed genes. Instead, we reasoned that an immune system response would be reflected by gene products that interact and are enriched for curated immunological functions or pathways. We found no interactions between the 27 genes that were enriched in the PROM cases genes using ingenuity pathway analysis (IPA), and these genes showed no statistically significant enrichment for immune response pathways. We interpreted this to mean that there is no consistent immune response in the cord blood of PROM cases. PROM was selected as a clinical event that incurred risk of infection for the foetus, but infection may not always occur. In addition, even where there is infection, there may be idiosyncratic responses reflecting heterogeneity in the type or timing of infection, or genetic or epigenetic variation in the individual host (5).

**Figure 1.**
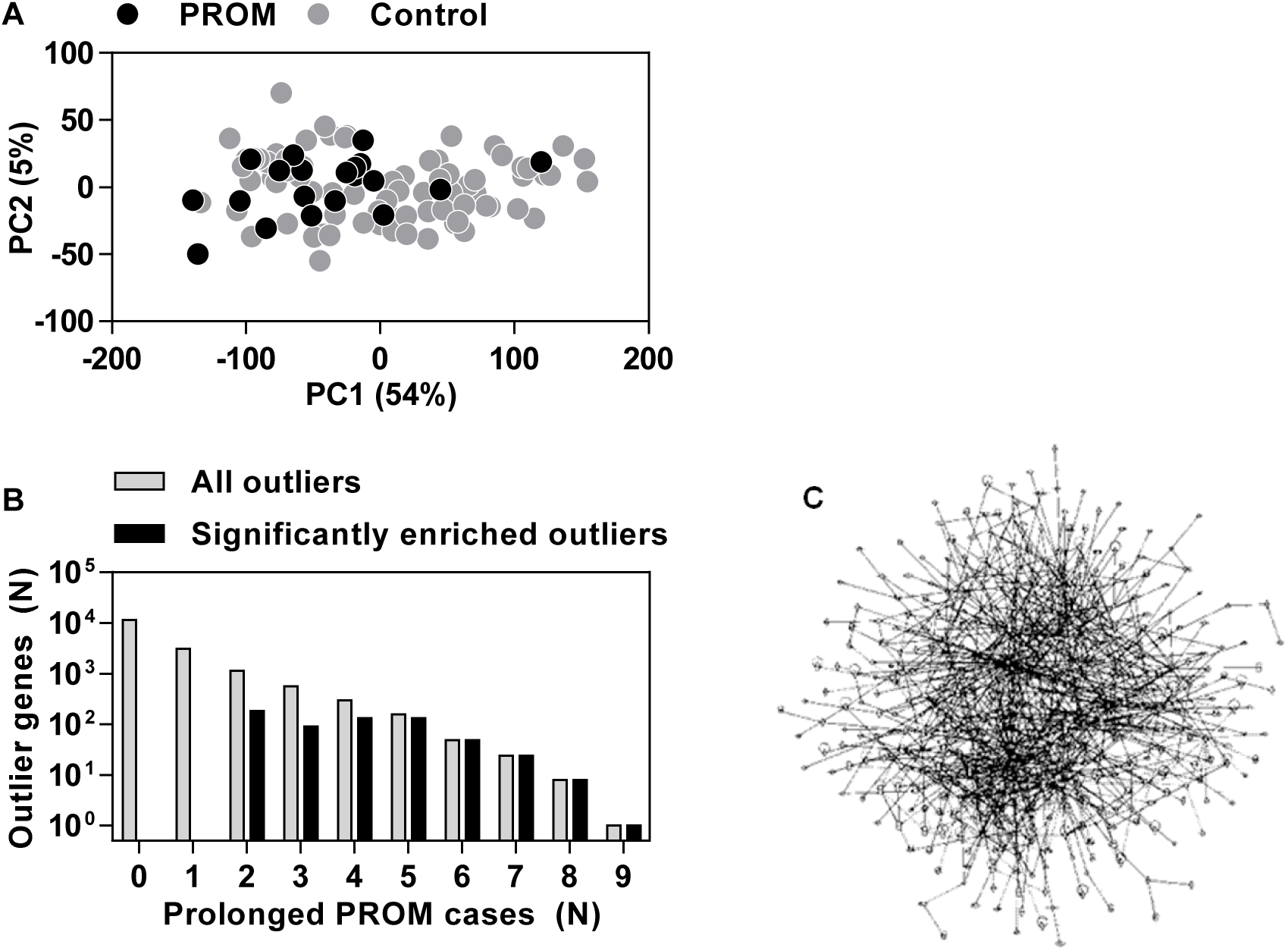
Identification of cord blood transcriptional changes associated with PROM. **(A)** Principle component analysis of cord blood transcriptomes with proportions of variance explained by each component given in brackets. **(B)** Total number of outlier transcripts expressed >1 Log_2_ higher in subsets of cord blood transcriptomes from prolonged PROM cases than the distribution of expression levels in controls (p<0.05, grey bars), and number of these outlier transcripts that were statistically enriched (Chi squared test, p<0.05, black bars) among PROM cases compared to controls. **(C)** Network of 381 directly and indirectly interacting genes among the statistically enriched outlier transcripts identified in (B).

In order to test the hypothesis that infection, and therefore an immune response, occurred in only a subset of PROM cases, we undertook outlier profile analysis (6) to identify transcripts in each individual PROM case that were significantly enriched compared to the controls, as a group (Figure 1B). This analysis would be expected to include outlier transcripts that may not be related to PROM and that are equally likely to occur in any control case compared to all others. Therefore, we focussed on only those outlier transcripts that were evident in a significantly higher proportion of PROM cases compared to controls. This analysis excluded all the outlier transcripts that were evident in only one PROM case, but identified 622 cord blood transcripts significantly over-represented in PROM cases (Figure 1B). Again to avoid type 2 errors we did not use multiple testing corrections in these analyses. Instead, we looked for evidence that outlier transcripts were statistically enriched for components of the immune system rather than being randomly selected from the genome. Although these transcripts represented the integrated list of outliers from different PROM cases, we reasoned that they encoded products with direct or indirect interactions within the immune system. To test this hypothesis, we first used IPA to identify the outlier genes among the PROM cases that are predicted to directly or indirectly interact with each other. This revealed one large network comprising 381 interacting genes (Figure 1C). We then evaluated the functional annotation of the interacting network of genes, using Reactome pathway enrichment analysis (Figure 2, Supplementary File 1). Immunological pathways revealed by this analysis were primarily those of innate immune signalling and antigen presentation, but not processes associated with adaptive T cell or B cell responses. On the basis that there has been no prior opportunity for priming of adaptive immune memory in the foetus, our results may reflect very early primary immune responses, in which adaptive immunity has not yet had time to emerge.

**Figure 2.**
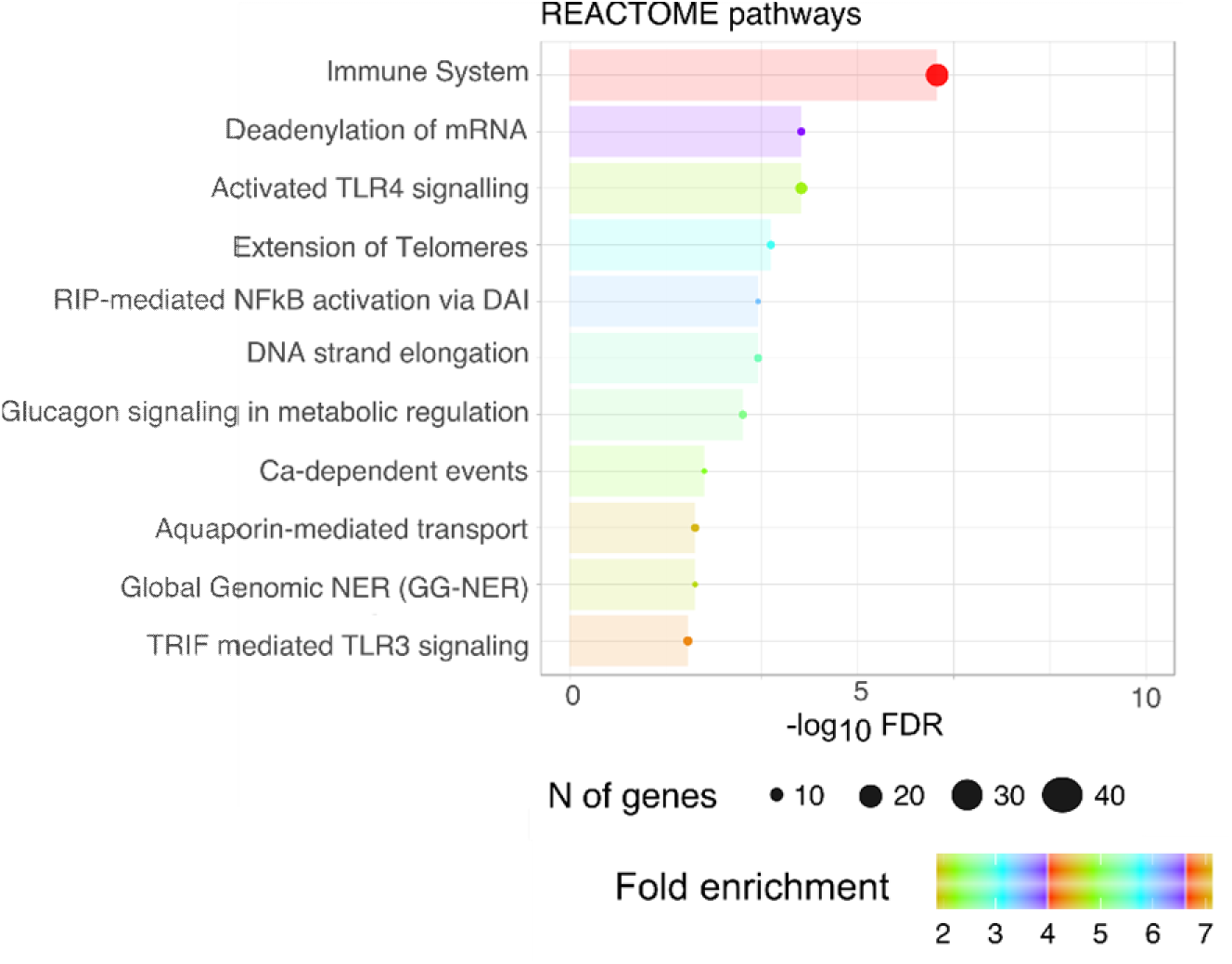
Functional annotation of the interactome of enriched transcripts in PROM cases. Summary of reactome pathways significantly enriched (FDR<0.05) amongst interactome of 381 outlier transcripts associated with PROM cases.

In order to confirm that these findings did not arise by chance, as a result of the very high dimensional data, we repeated the same analysis, using 100 iterations of 20 cases subsampled from the complete data set. We identified outlier transcripts that were significantly enriched in the subsample on each iteration (Supplementary Figure 1). These occurred at substantially lower frequency (median of 100 outliers per iteration, 95% confidence interval 69-140) compared to the 622 outlier transcripts identified in PROM cases. This analysis confirmed that the number of outlier transcripts among the PROM cases was significantly greater than would be expected by chance. Next we sought to test whether immune system pathway enrichment was specific to the integrated list of PROM outlier transcripts. To do this we collated all the outlier transcripts from the 100 iterations of 20 random cases subsampled from the entire data set, excluding rare outliers that occurred in less than 5% of iterations. This provided a list of 707 outlier transcripts which occur with >95% confidence in random sampling. Interactome analysis suggested that 185 of these transcripts may participate in a functional network. Pathway analysis of this interactome revealed significant enrichment for several non-immune processes, but even these were only represented by very few genes (Supplementary Figure 1, Supplementary File 2). Taken together, these data show that enrichment of transcripts associated with innate immune responses was specific to a subset of PROM cases.

Finally, we tested the hypothesis that the heterogeneity of transcriptional changes in subsets of PROM cases was due to idiosyncratic responses. We undertook hierarchical clustering of the PROM cases on the basis of the average Jaccard distance of outlier transcripts, and likewise clustered the 381 interacting outlier genes on the basis of the average Jaccard distance of the PROM cases in which they were identified (Figure 3). In this analysis, the clustering of cases was associated with accumulation of outlier transcripts *per se*, rather than outlier transcriptional profiles composed of selected combinations of genes. We infer from this that transcriptional changes in PROM are mostly idiosyncratic, and that the cumulative number of changes, rather than the precise nature of changes, may provide a surrogate for the extent of the immune response. Of note, in this sample, variation in immunoreactivity (as defined by the frequency of outlier transcripts in each case) did not correlate with variation in any of the variables in the clinical metadata (Table 1). Much larger sample sizes will be needed to identify the clinical correlates of different in immune signatures in cord blood.

**Figure 3.**
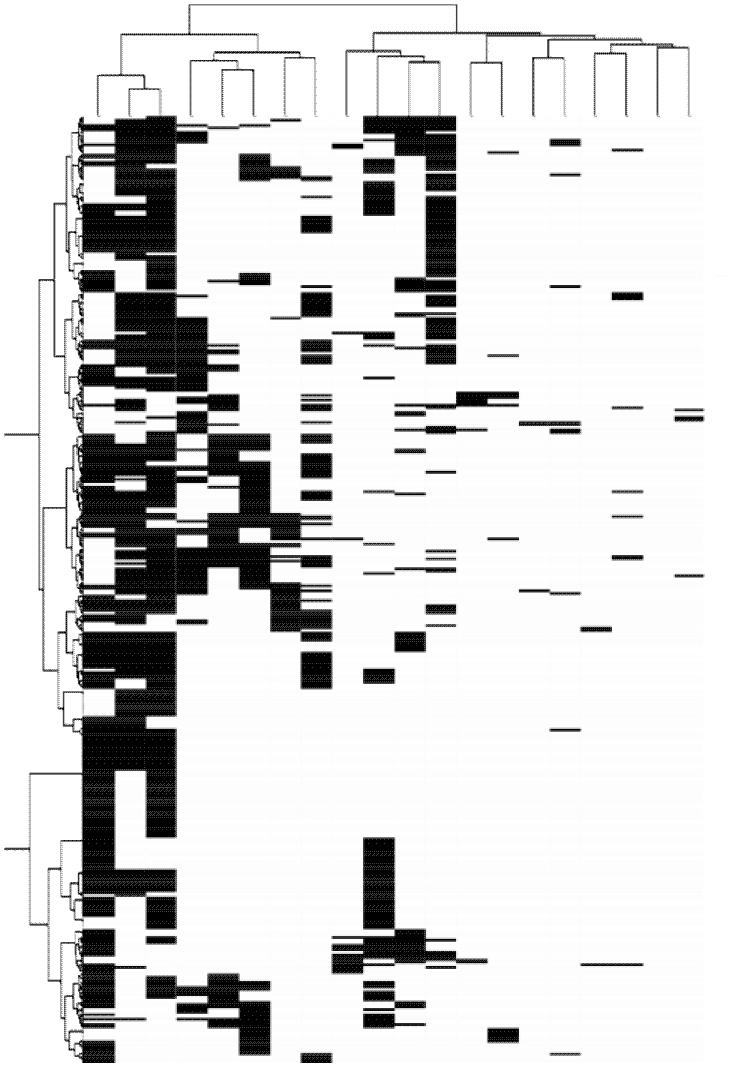
Clustering of cases and outlier transcripts among PROM cases. Hierarchical clustering of cases (columns) and outlier (transcripts) by average Jaccard distance.

We conclude that transcriptional profiling of cord blood can be used to identify foetal immune responses at the systems level and with molecular resolution. The present study provides the first direct evidence that a subset of PROM cases is associated with foetal innate immune responses, strongly supporting the prevailing view that PROM may increase the risk of perinatal infection. These findings pave the way for larger studies incorporating longitudinal outcome data to correlate transcriptional changes in the cord blood with clinical outcomes in infancy and childhood, and thereby identify biomarkers to stratify clinical risk to support precision targeting of interventions such as antimicrobial therapy and enhanced monitoring. In addition, our findings encourage the application of cord blood transcriptomics to identify immunological and inflammatory pathways that may be used to stratify clinical risk and offer novel targets for host-directed therapies in other examples of congenital infection. By scaling up this approach to provide adequate statistical power, transcriptional profiling of cord blood may also provide the opportunity to investigate the impact of common genetic variation and other prenatal exposures on innate immune responses in the neonate, which may influence immunological health in later life.

## Methods

### Study approval

This study was approved by the UK National Research Ethics Service (reference: 12/LO/1492).

### Study population and sampling

Pregnant women were recruited to the study at the time of antenatal visits to hospital. All participants provided written informed consent. Cord blood samples were collected directly into Tempus^TM^ tubes immediately after delivery. Selected demographic, clinical laboratory and clinical outcome data were obtained from the hospital electronic data repository and medical notes. PROM cases were defined as delivery more than 24 hours after rupture of membranes.

### Cord blood transcriptional profiling

RNA was extracted using the Tempus™ Spin RNA Isolation kit (Applied Biosystems) or PAXgene 96 Blood RNA Kit (PreAnalytiX). Genomic DNA was removed with the TURBO DNA-free™ kit (Ambion). RNeasy MinElute Cleanup kit (Qiagen) was used to concentrate the RNA before globin mRNA depletion with GLOBINclear™ kit (Ambion) and RNA quality control was assessed using the Agilent 2100 Bioanalyzer (Agilent Technologies). Fluorophore labelled cRNA was then generated using the Low Input Quick Amp labelling kit, and hybridised to SurePrint G3 Human Gene Expression v3 8×60K or Human Gene Expression v2 4×44K whole genome microarrays (Agilent Technologies). Array images were acquired with Agilent’s dual-laser microarray scanner G2565BA and analysed with Agilent Feature Extraction software (v9.5.1). Log_2_ transformed median Cy3 and Cy5 signal intensities were normalized using LOESS local linear regression against the mean signal of all the samples using the R package agilp (7, 8). All microarray data used in this study are available in ArrayExpress (https://www.ebi.ac.uk/arrayexpress/) under the accession number E-MTAB-6431.

### Data analysis

Analysis of microarray data was conducted on log_2_ transformed data (8), and restricted to probes expressed above background negative control levels in at least one sample. Principal component analysis was conducted using the prcomp package in the R statistical computing platform. Significant gene expression differences for protein coding genes with >1 log_2_ difference between data sets were identified using Mann Whitney tests for non-parametric data in MultiExperiment Viewer v4.9 (http://www.tm4.org), with p value<0.05. The Zodet package in R statistical was used to perform outlier profile analysis based on (8), using a p value<0.05 with a fold difference filter of >1 log_2_. Chi squared tests implemented in R were then used to identify outlier transcripts statistically enriched in cases compared to controls or in subsamples of control cases, using a p value<0.05. Interactomes were identified using Ingenuity Pathway analysis (www.ingenuity.com) and Reactome pathway enrichment analysis were performed in XGR (9), filtered from pathways in the Reactome database with more than 10 components represented by at least 3 genes in the list of genes being analysed and enrichment with a false discovery rate (FDR)<0.05. K-means clustering of Jaccard indices to quantify the similarity between the composition of genes mapping to each pathway was used to identify a maximum of 10 groups of pathways, from which the pathway consisting the largest total number of genes was selected to provide a representative annotation for each group. The data were plotted using the ggplot R package and graphpad prism.

## Author contributions

AR, ER, NF, PB and MN conceived of the study. ET, SB, AS and NS coordinated sample and clinical metadata collection. ET, CT, AS and CK undertook sample processing. AL, BMC, ET, CT, CV and MN undertook data analysis. ET, CT and MN wrote the manuscript with input from all the authors.

## Acknowledgments

We thank the study participants as well as the research midwife teams and hospital staff at University College London Hospitals and Barking, Havering and Redbridge University Hospitals NHS Trusts, that supported the recruitment of participants. This study was funded by the Wellcome Trust (WT101169MA) and by the National Institute for Health Research Biomedical Research Centre at University College London Hospitals. NF was funded by a National Institute for Health Research (NIHR) Academic Clinical Lectureship.

---

**Supplementary Figure 1.**
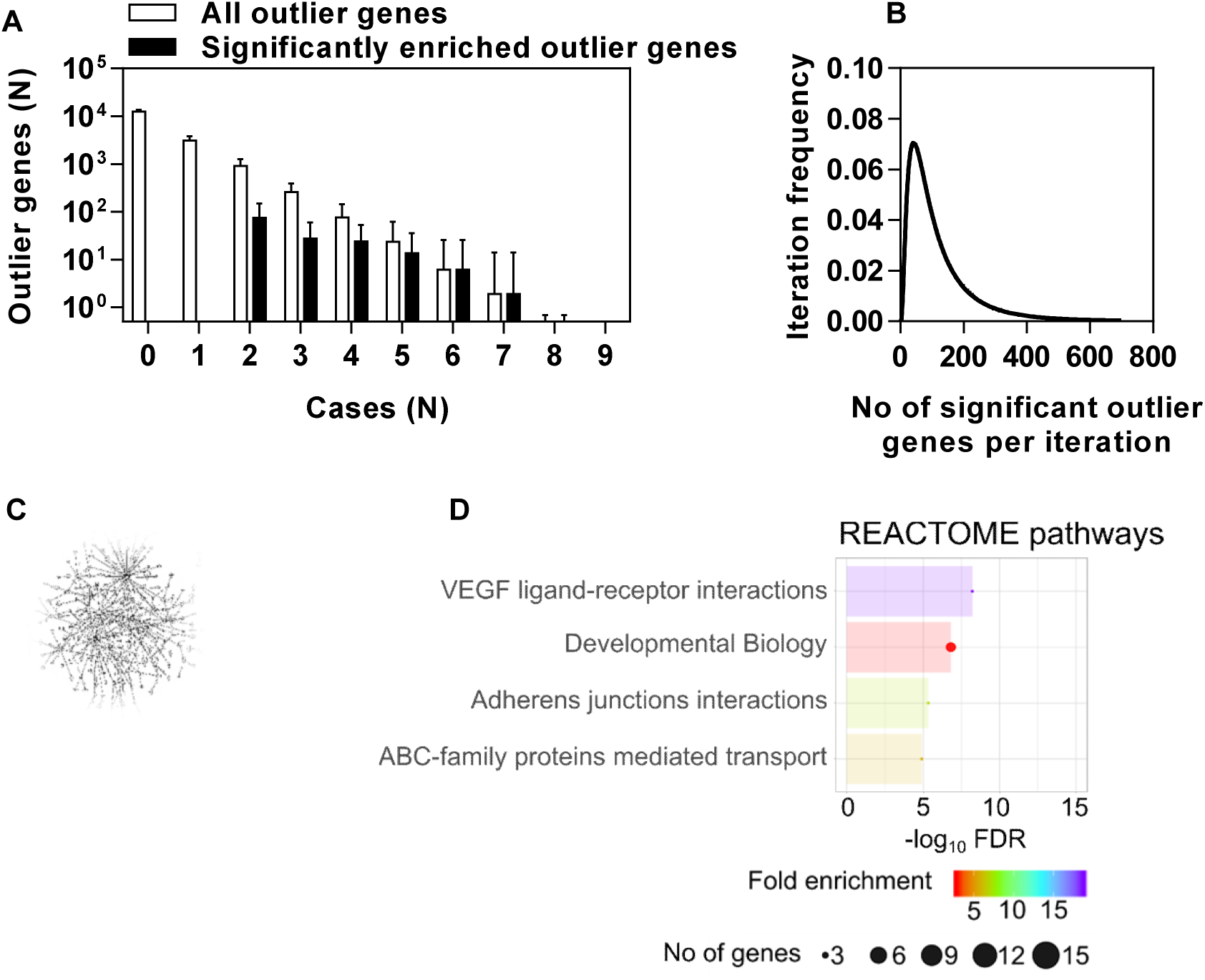
Cord blood outlier profile analysis amongst control cases. **(A)** Total number of outlier transcripts in multiple subsamples of 20 cases from all PROM and control cases (expressed >1 Log_2_ higher levels than the distribution of expression levels of the remaining cases in each iteration, p<0.05 grey bars) and number of these outlier transcripts that were statistically enriched (Chi squared test, p<0.05, black bars) among subsamples compared to remaining controls in each iteration (bars represent mean±SD). **(B)** Frequency distribution of the number of significant outlier transcripts in 100 subsamples from the control cases. **(C)** Network of 185 directly and indirectly interacting genes among the integrated list of statistically enriched outlier transcripts identified in ≥5% of subsamples in (A-B). **(D)** Summary of reactome pathways significantly enriched (FDR<0.05) amongst interactome of 185 outlier transcripts in (C).

## Notes

**Conflict of interest statement** The authors have declared that no conflict of interest exists.

